# *Operando* Monitoring of Metabolic Fluxes In Cholestatic Murine Liver Tissue Slices By Microfluidic NMR

**DOI:** 10.1101/2025.10.08.681085

**Authors:** Sylwia J. Barker, Bishnubrata Patra, Manvendra Sharma, Ruby E. H. Karsten, Elisabeth Verpoorte, Marcel Utz

**Affiliations:** School of Chemistry, University of Southampton, UK; Faculty of Science and Engineering, Pharmaceutical Analysis, University of Groningen, Netherlands; Pharmaceutical Analysis, Groningen Research Institute of Pharmacy, University of Groningen, Netherlands

## Abstract

Monitoring metabolism in living tissues with high temporal resolution and broad metabolite coverage remains a major challenge. We introduce TISuMR, a microfluidic Lab-on-a-Chip platform that enables continuous, non-invasive *operando* NMR spectroscopy of live tissue slices. The TISuMR platform replaces the conventional NMR sample tube with a fully integrated microfluidic culture system that maintains tissue viability through dynamic nutrient perifusion, gas exchange via a diffusion membrane, and precise temperature control. Coupled with a custom-designed micro-NMR probe, the platform allows detection of nearly two dozens of metabolites from just 2.5 *µ*L of sample. In a proof-of-concept study, we demonstrate the platform’s unique ability to resolve dynamic metabolic fluxes and to monitor, in real time, the onset of chlorpromazine-induced cholestasis in murine liver tissue, with a time resolution of just over three minutes. This approach provides a powerful, minimally disruptive tool for studying tissue metabolism in real time.

## Introduction

Metabolic changes are a crucial hallmark of cellular health, as cells modulate their metabolism in response to different physiological and pathological conditions.^1–7^ Therefore, continuous real-time monitoring of metabolic activity in living biological systems can enhance our understanding of their function and the mechanisms behind disease and toxicity. While animal models remain important, progress in the life sciences increasingly relies on cultured model systems, such as tissues, cell aggregates, and individual cells, which are supplied with nutrients and oxygen in a controlled manner. Such model systems avoid many of the ethical concerns associated with animal models, and also have the potential to provide data that is more relevant to human disease, as human cells can be used directly. They also provide more controlled conditions, allowing the consistent repetition of experiments, which facilitates the derivation of statistically relevant results. Last, but not least, such models provide much easier access for analytical methods of various kinds, and thus can provide a rich set of data on morphology (microscopy), expression levels of surface markers (fluorescence-activated cell sorting) and enzymes (qPCR-based transcriptomics), on genotype and mutation (genomics), and, finally, on the metabolic activity of the cells. As the metabolism lies at the top of the cascade of biological activity that is anchored in the genome, it tends to exhibit the most immediate response to changes in the cellular environment. Quantification of the fluxes in the network of metabolic reactions therefore provides a window into the inner workings of biological systems. Culture assays often benefit from microfluidic lab-on-a-chip (LoC) devices, which provide a convenient integrated platform for culture and downstream analytics.

While destructive end-point analysis of culture assays is well established, non-invasive quantification of the metabolic activity could provide valuable, complementary information. This has sparked interest in *in-situ* sensing technologies. These are often based on electrochemical detectors, which offer high sensitivity, rapid response times, and ease of integration. However, they can only target a narrow spectrum of metabolites, and studies are often focused on a single analyte like glucose, lactate, pH, or dissolved oxygen.^8–11^ This selectivity is inherent to the design of electrochemical sensors, which detect specific redox-active species. Multiple metabolites require an array of such sensors, increasing the complexity of the setup. On the other hand, mass spectrometry (MS) can identify a wide range of metabolites, particularly if combined with chromatographic separation,^12^ but generally requires terminating the experiment and destroying the sample. This invasive nature precludes real-time monitoring of metabolic changes and limits the ability to observe dynamic biological processes as they occur. In contrast, high-resolution Nuclear Magnetic Resonance (NMR) spectroscopy allows non-invasive quantification of biochemical processes and can provide detailed information about the metabolic state of biological systems without disturbing the biological function of the cultured cells. Seamless integration of NMR with culturing platforms is challenging, in particular in cases where the biological system to be observed is small. Conventional liquid-state NMR experiments are carried out in 5 mm glass tubes, and require relatively large sample volumes (~ 500 − 600 *µ*L). Although *in situ* experiments with cells cultured in an NMR tube have been demonstrated, they require a large number of cells (*>* 10^9^), and offer only limited options to provide physiologically relevant conditions.^13^

Microfluidic Lab-on-a-Chip (LoC) technology provides a versatile and efficient platform for culturing biological systems in highly controlled microenvironments.^14–17^ These devices enable precise manipulation of fluid dynamics, nutrient delivery, and waste removal at the microscale to provide more relevant physiological conditions compared to traditional static cell culture systems. LoC platforms can integrate experimental functions such as cell culture, analysis, and monitoring, into a single device. The miniaturized scale reduces the consumption of reagents and sample, making them ideal for experiments with limited biological materials. Integrating LoC technology with NMR spectroscopy is challenging as on the one hand, the microstructure of the device tends to interfere with the homogeneity of the magnetic field, and, on the other hand, the small volume places high demands on sensitivity. It has been shown that NMR detection sensitivity scales favourably if the sample and the detector are miniaturized at the same time.^18–21^ Magnetic field homogeneity can be maintained by careful selection of materials used in microfluidic chips,^21–24^ and by optimizing the microstructure of the device such as to minimize susceptibility effects in the sampling volume.^25,26^ The development of planar NMR detectors^24^ that can directly accommodate microfluidic devices^22,27^ has enabled quantification of metabolic turnover of adherent cells and cell spheroids cultured in LoC.^28,29^ Rogers *et al*.^30^ demonstrated that time-resolved metabolic fluxes can be determined from as few as 500 cells cultured within microfluidic devices over a 24 hour period.

Culture models based on a single cell type fail to capture the intercellular crosstalk and extracellular matrix-mediated modulation, which can be critical for accurately modeling disease processes.^31,32^ *Ex vivo* tissue-slice cultures, as an intermediate between traditional cell cultures and *in vivo* animal models, offer a more physiologically relevant platform that preserves the structural and cellular heterogeneity of native tissues. Tissue-specific processes can thus be studied in a controlled *ex vivo* environment that preserves the diversity of cell types and their extracellular context, including the native network of cell-cell contacts.^33^ The literature documents methods for the culture of a wide range of tissue types,^34–40^ underscoring the growing utility and adaptability of this approach to study many diseases, for example fibrosis^41^ and cancer^42^ of the liver, and infectious diseases of the lung.^36,37^

Magnetic resonance (MR) experiments have also been applied to the study of tissue-slice culture. Shepherd *et al*. (2006)^43^ obtained MR images of rat hippocampal slices using a 10 mm NMR tube. Harris *et al*.^44^ reported a method to continuously monitor the metabolism of *ex vivo* brain slices with a time resolution of 1-2 minutes using hyperpolarized 1−^13^C[pyruvate] ^31^P NMR. Hyperpolarization enhances the MR signal by several orders of magnitude by increasing the population difference between nuclear spin states above its thermal equilibrium value. This allows for the quantification of metabolites and dynamic processes below the detection threshold of conventional MR. Hyperpolarized [1−^13^C]pyruvate was also used to study precision-cut tumor slices of breast cancer xenografts,^45^ liver slices,^46,47^ brain slices^48–50^ as well as an *ex vivo* perfused mouse heart (the Langendorff heart).^51 31^P NMR has also been used to non-invasively monitor tissue-slice viability through the ATP signal,^44^ and to quantify the extracellular volume using the biologically inert tracer 3-aminoproponyl phosphate.^52^ These studies were carried out in conventional NMR tubes that were filled with multiple tissue-slices. This does not allow precise control over conditions such as oxygenation, nutrient delivery, and waste removal. Additionally, the large volume of culture media required in NMR tubes can dilute metabolite signals, reducing sensitivity and necessitating higher concentrations of hyperpolarized substrates, which may not be physiologically relevant.

In the following, we present a microfluidic culture and analytical platform that enables real-time metabolomic studies of living tissue slices *in operando* through high-resolution NMR spectroscopy. The platform comprises a microfluidic chip to culture the tissue-slice and a fluidic interface for provision of nutrients and control of physiological conditions. The interface connects the chip to the external gas and nutrient supplies as well as temperature control, which are necessary to maintain the viability of tissue inside an NMR spectrometer. Using our platform, we demonstrate the ability to simultaneously observe the production of dozens of metabolites from a murine liver-tissue slice. Lastly, we show that our platform can be applied to study a disease model by introducing chlorpromazine into the system, a drug known to induce liver cholestasis.

## The TISuMR platform

The TISuMR microfluidic device comprises a central culturing chamber, designed to house a tissue-slice, surrounded by fluidic channels that enable delivery of oxygen and nutrients to maintain the viability of the tissue, as shown in Fig.1b. Metabolites are detected in the 2.5 *µ*L NMR detection chamber located in the lower part of the device. The operating principle behind the device is illustrated in Fig.1c-d. Initially, a 25 *µ*L plug of medium is introduced into the liquid channel, marked in pink, using a unidirectional pump. Carbogen gas (95% O_2_ and 5% CO_2_)^53,54^ flows through the gas channel (blue). Its diffusion into the liquid channel is facilitated by a poly(dimethylsiloxane) (PDMS) membrane, which acts as a bridge between the two channels as shown in Fig. 1c. At 25 °C and 1.0 atm, solubility in water is 1.22 mM.^55^ To ensure oxygen saturation of the medium, the device was designed with an ex-tended path length to enhance the surface area available for diffusion. Oxygen uptake in this type of geometry has been quantified by Yilmaz et al,^56^ who reported a characteristic time of 0.42 s for equilibration of the oxygen partial pressure between the liquid and gas channels. At the flow rate used here (about 8 *µ*Lmin^−1^), the medium spends about 12 s in contact with the PDMS membrane, more than 20x the equilibration time. A similar approach, has been used successfully for biphasic gas-liquid hydrogenation reactions on a chip.^57–59^ The side-by-side configuration prevents compressing the membrane into the channels, and enables the use of elevated pressures enhancing gas dissolution efficiency.

**Figure 1:**
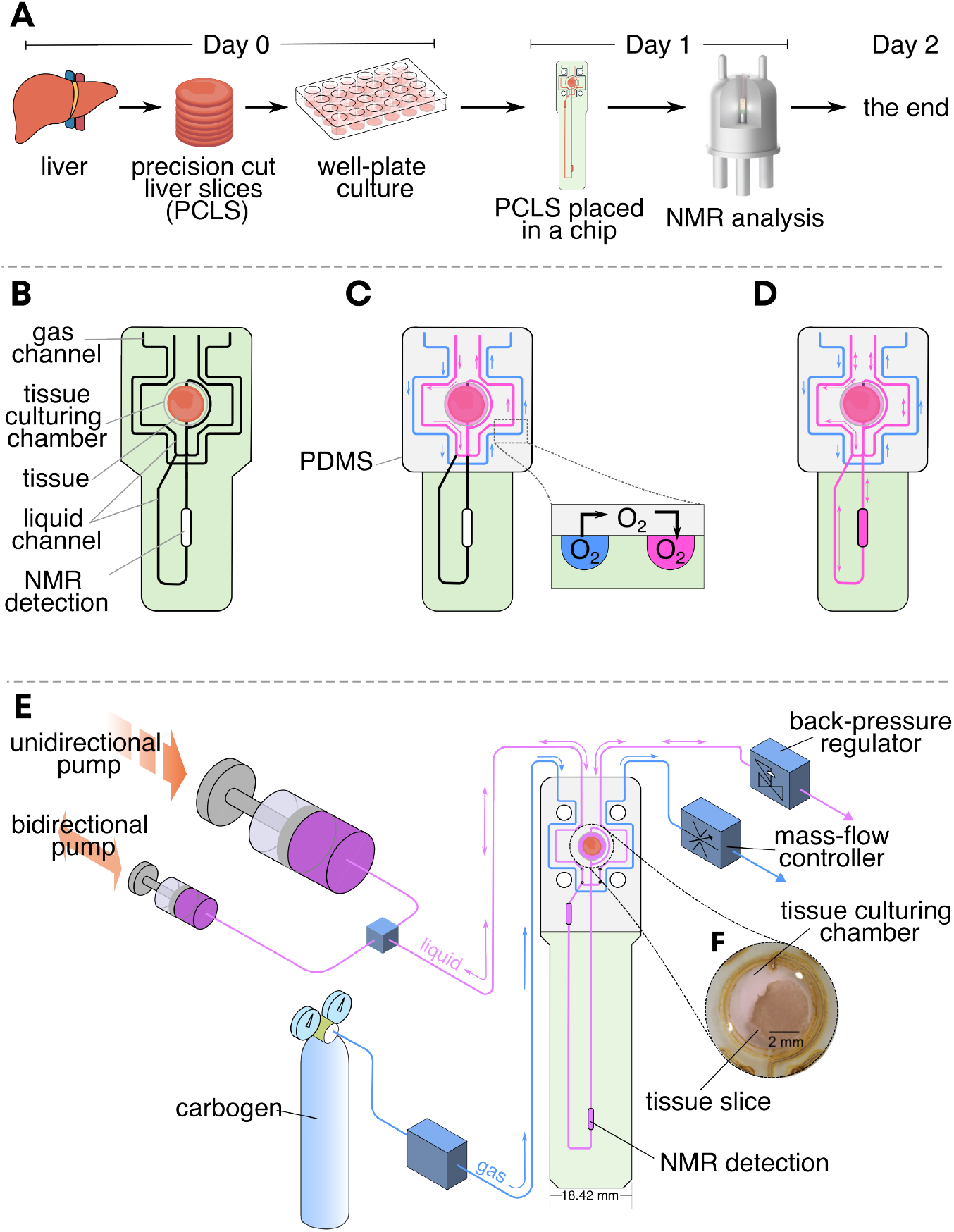
a) Experimental timeline. On Day 0 mouse liver is cut into slices and placed in a well plate for overnight culture in a tissue culture incubator. On Day 1, PCLS is placed into a micorfluidic device and observed in an NMR spectrometer. b-d) Operating principle behind the TISuMR plarform. e) Experimental set up used for maintaining viability of the PCLS in a microfluidic chip. f) A microscope image of the tissue culturing chamber.

As a proof-of-concept, we utilized liver tissue from C57BL*/*6 mice, leveraging tissue availability from ongoing studies at a collaborating institute. This approach maximized experimental output from a single-animal sacrifice, aligning with the principles of the 3R (Replacement, Reduction, and Refinement) initiative^60^ and underscoring the potential of microfluidic technology to advance ethical and sustainable research practices. Precision-cut liver slices (PCLS) were prepared following a standard protocol.^53,54^ The key steps are illustrated in Fig. 1a. Briefly, liver cores were prepared using a 6-mm biopsy punch and sliced into PCLS using a Krumdieck Tissue Slicer (Alabama Research and Development, USA). The resulting 250-*µ*m-thick PCLS were incubated overnight in a standard 12 well-plate at 37°C and an atmosphere of 80% oxygen and 5% CO_2_. On Day 1, one PCLS was used for NMR analysis to assess metabolic activity. The perifusion flow rate is set by the rate of oxygen uptake of the tissue and the oxygen solubility of the medium. As discussed in detail below, unidirectional flow at this rate did not result in concentration changes in the flowing media that could be detected by NMR. This illustrates a fundamental dilemma that all attempts to quantify metabolic fluxes through concentration measurements must face: The biological system needs to be kept in homeostatic conditions to ensure viability. However, for quantification to be possible, significant nutrient depletion and accumulation of physiological products must occur. In the present work, we have solved this problem by passing a plug of culture medium of 25 *µ*L volume back and forth over the tissue. The chip is designed in such a way that the plug comes in contact with a gas exchanger on each pass, thus ensuring that the oxygen- and CO_2_ concentrations of the fluid arriving at the tissue are always in equilibrium with the supplied gas (carbogen). To this effect, a Y-junction was incorporated to connect a bidirectional pump, as shown in Fig. 1e. After introducing the medium plug, the bidirectional pump facilitated a back-and-forth (BNF) flow of the medium at a rate of 8 *µ*L min over a duration of 3.5 hours. This ensured effective perifusion and observation of metabolic changes. After that time, a fresh plug of medium was introduced to replenish the nutrients. This cycle was repeated for the entire time that the PCLS was inside the NMR spectrometer. The outflow of the chip was connected to a back-pressure regulator, maintaing a positive pressure of 1.2 bar with respect to ambient; this was found crucial in order to maintain the system free of bubbles.

Fig. 2a depicts the assembly used to maintain the viability and perifuse the tissue inside of an NMR spectrometer. It consists of the microfluidic device interposed between two PDMS membranes, all held securely by a fluidic interface. The front PDMS membrane covers the gas and liquid channels, enabling gas exchange to oxygenate the liquid channel, while the rear PDMS membrane aids sealing to ensure leak-proof operation. Additionally, the fluidic interface is equipped with fluidic connectors that align with the inlets and outlets on the device. These connectors enable seamless integration with external components, such as pumps for nutrient and media delivery and a gas cylinder for supplying the required gas mixture. To maintain the physiological temperature of 37°C required for optimal tissue function, the fluidic interface was equipped with a precision PID-controlled heating system, as described in detail in.^30^ This system continuously monitors and regulates the temperature of the device, ensuring uniform thermal conditions throughout the duration of the experiment. The PID controller operates by dynamically adjusting the heating output based on real-time feedback from temperature sensors embedded within the fluidic interface. NMR spectra are acquired at the 2.5 *µ*L NMR sample detection chamber using a custom-built probe equipped with stripline-based micro-NMR detector.^22^ In this way, the viability of a liver tissue slice was maintained inside of the an NMR spectrometer while continuously monitoring its metabolic output (Fig. 2b).

**Figure 2:**
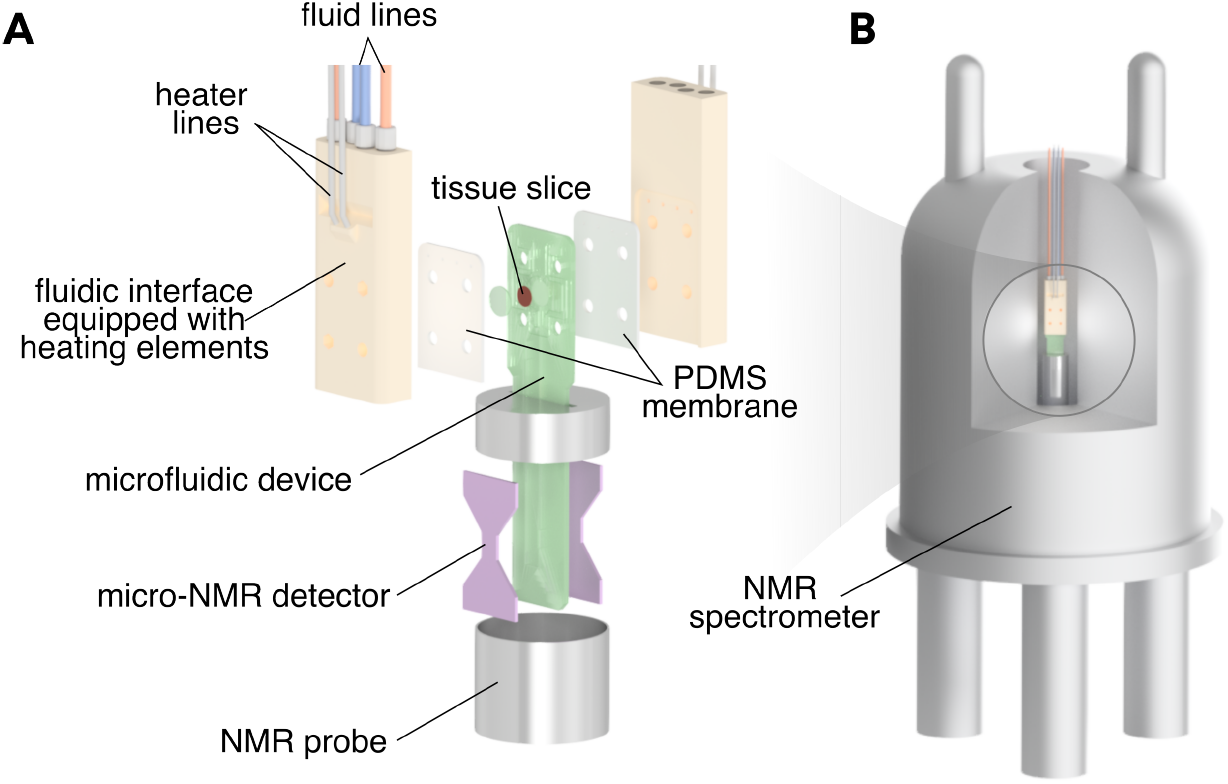
a) A rendering of the TISuMR assembly. The microfluidic device and PDMS membranes were secured together by the fluidic interface. This interface was connected to an external supply of nutrients and oxygen, enabling us to maintain appropriate culturing conditions inside of the NMR spectrometer. Additionally, the interface functioned as a heater, providing a stable temperature of 37 °C. b) The entire microfluidic assembly was placed in an NMR spectrometer for 24 hour observation.

## Results and Discussion

A typical NMR spectrum obtained from the 2.5 *µ*L sample detection chamber in a 14.1 T spectrometer is shown in the top trace (blue) of Fig. 3. The spectra reported here are an average of 48 scans obtained every ~ 3 minutes. After apodization with 1 Hz Lorentzian line broadening, the signal-to-noise ratio (SNR) of the spectrum was determined using the DMSO peak, resulting in a value of 506. The orange trace shows the fitted spectrum obtained through spectral decomposition using reference peak positions from the Human Metabolome Database,^61^ following a protocol reported previously.^30^ The bottom panel showcases the 11 individual metabolites and DMSO identified within the spectrum. The fit closely aligns with the experimental spectrum, quantitatively accounting for most of the visible signals. Exceptions include peaks marked with an asterisk at 1.2, 2.95 and 3.2 pmm. The peak at 1.2 ppm is attributed to residual isopropanol from the fabrication process, while the other two peaks correspond to exchangeable protons in the deuterated HEPES buffer.^13,30^

**Figure 3:**
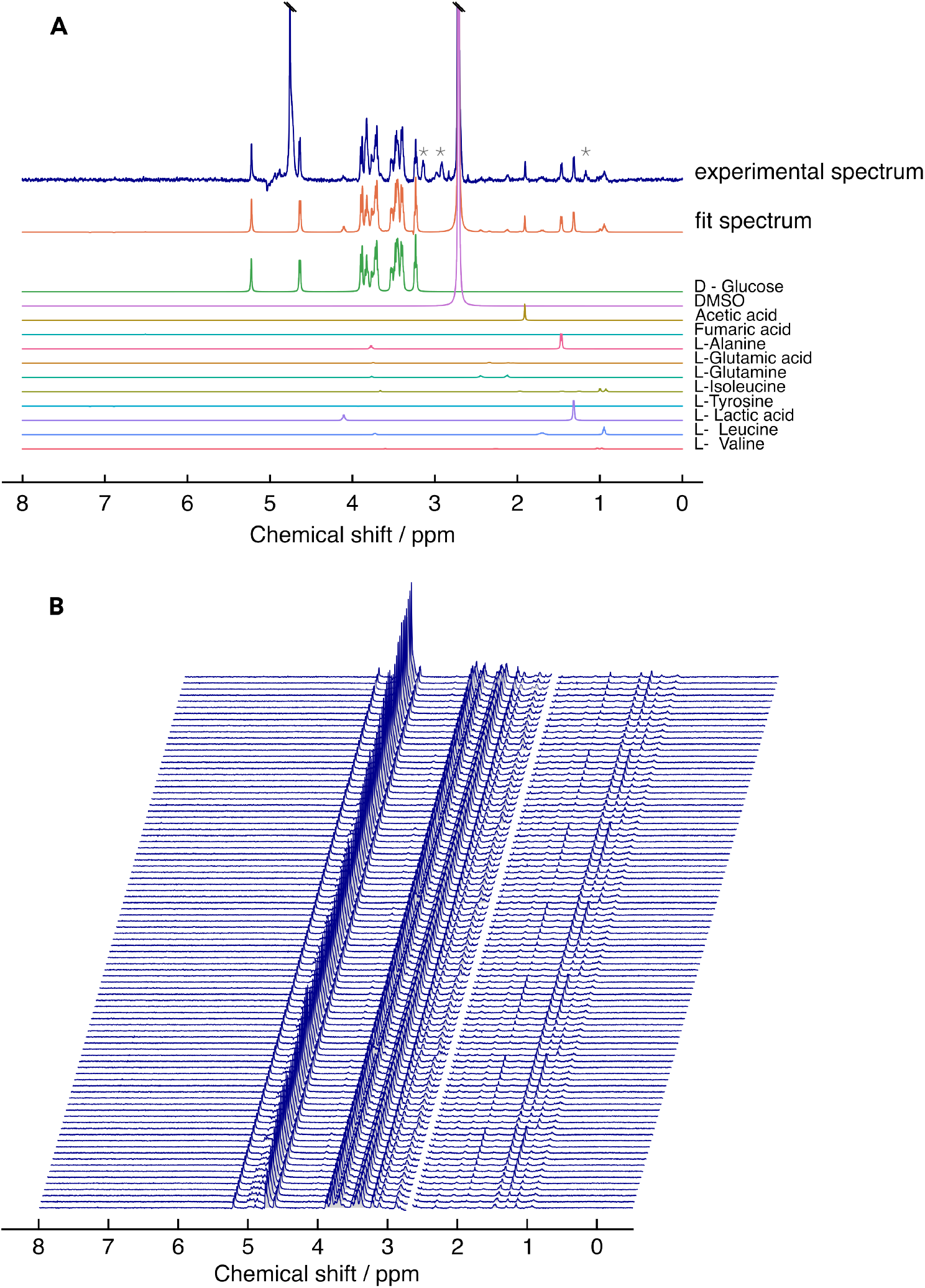
a) Example of an experimental spectrum and its decomposition into constituent metabolites. Peaks marked with an asterisk correspond to residual isopropanol (1.2 ppm) from the fabrication process, and HEPES. b) Sequential NMR spectra recorded throughout the experiment, displayed in a waterfall plot. DMSO peak has been suppressed for spectral clarity.

Fig. 3b illustrates the continuous data acquisition process, and the consistency of the data over 24h highlighs the robustness and efficiency of the microfluidic platform for long-term metabolomic studies. Absolute concentrations were determined by using the known concentration of glucose (25 mM) in the first spectrum as the reference standard for calibration. This initial value was used to scale the signal intensities of all detected metabolites, allowing for the quantification of their concentrations across the entire dataset.

Fig. 4 shows the time-resolved concentration profile of 11 metabolites and DMSO obtained from a single 24 hour observation of a precision-cut liver tissue slice cultured within an NMR spectrometer. The blue traces depict experimental measurements, while the red lines represent separate linear fits to the data. A fresh plug of medium was used to perifuse the tissue every 3.5 hours, resulting in a distinctive pattern observed in the concentration profiles. For example, the concentration of acetic acid increases from 0 to 1.2 mM over the first 3.5h period (Fig. 4B). As the first plug is replaced with fresh medium, the concentration returns to the initial value of 0, and then resumes the upward trend. A similar sawtooth pattern is exhibited by most metabolites to different degrees. While the concentrations obtained from individual spectra exhibit fluctuations due to the spectral noise, linear fits are very consistent for most metabolites. The fits shown as red lines in Fig. 4 are given by

**Figure 4:**
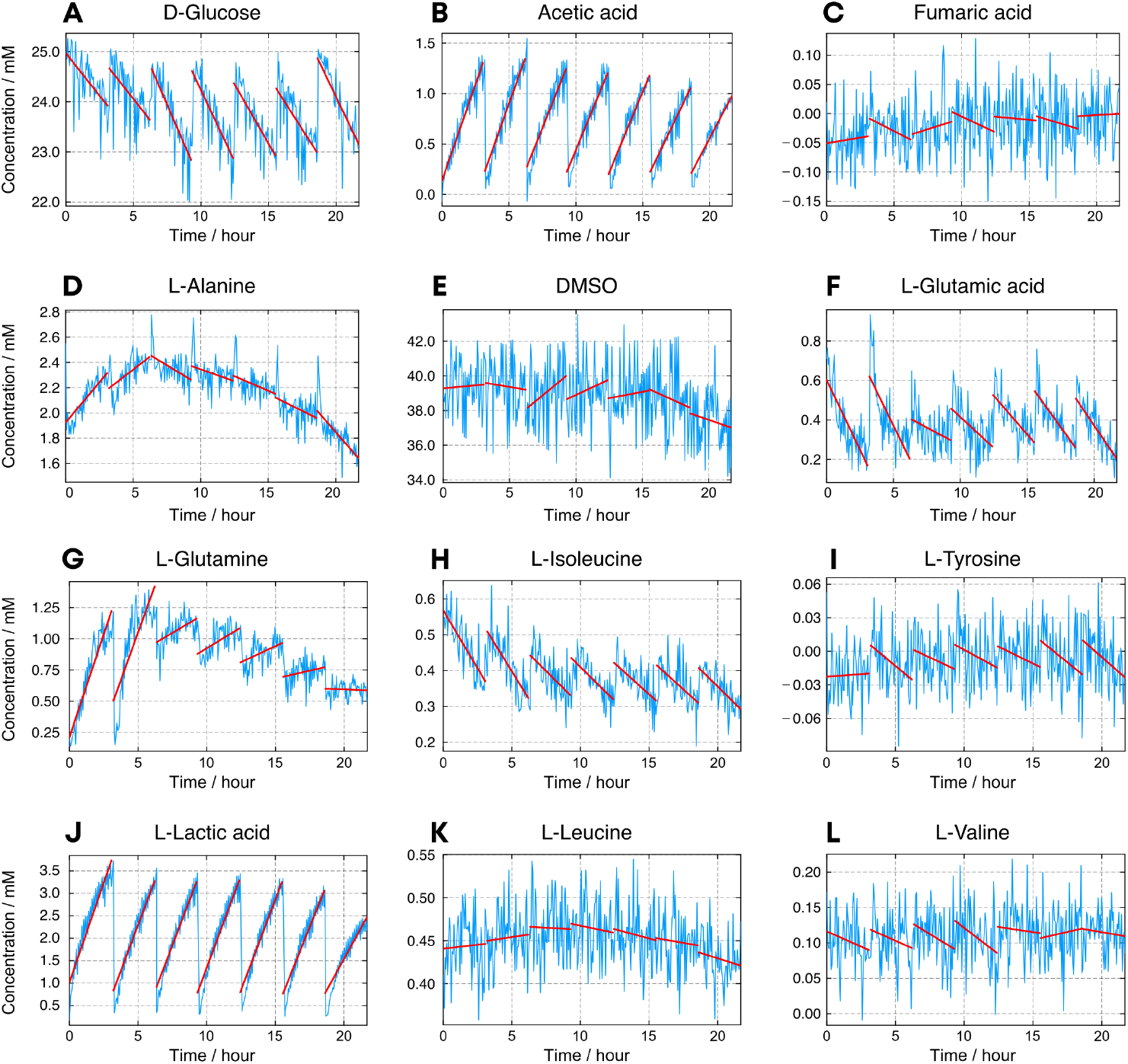
a-l) Time-resolved metabolic profiles of metabolites identified during PCLS culture on the TISuMR platform. The blue curves are obtained from individual NMR spectra, while the red lines represent linear fits to the data.

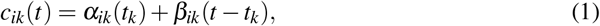

where *i* runs over all metabolites, 0 < *k* ≤ 7 runs over the intervals, and *t*_*k*_ are the interval starting times. The starting concentrations (*α*–*ik*) as well as the rates (*β* –*ik*) have been determined by linear regression. Notably, the concentration of D-glucose (a) decreases gradually over time, reflecting its steady consumption by the tissue. We observed an increase in the glucose consumption rate over the experimental period. Initially, the rate was found to be −0.340 (90%CI: 0.191,0.106)mMh^−1^, which nearly doubled to −0.623 (90%CI: 0.192,0.115)mMh^−1^ in the third period. L-lactic acid displays a different pattern with an initial production rate of +0.886 (90%CI: 0.145,0.081)mMh^−1^ that slowly decreases over time. In the last period, the lactic acid production was found to be +0.55 (90%CI: 0.196,0.109)mMh^−1^. Rates of other metabolites such as L-tyrosine (Fig. 4i), L-leucine (Fig. 4k) and L-valine (Fig. 4l) remain stable over the observation time.

Figure 5 shows the time-resolved metabolic profiles of L-lactic acid, acetic acid, L-glutamine, and L-tyrosine, along with their corresponding metabolic rates (*β*_*ik*_). L-lactic acid exhibits consistently high production rates (~ 0.75 – 0.9 mMh^−1^), with a slight decline toward the later time points. Acetic acid shows a gradual decrease in production rate from (~ 0.4 to ~ 0.25 mMh^−1^ over the 24-hour period. L-glutamine shows a high initial production rate (~ 0.3 mMh^−1^), which decreases sharply and stabilizes around (~ 0.05 mMh^−1^), suggesting an early metabolic response followed by reduced demand or entry into a steady state. Lastly, data from individual spectra of L-tyrosine is too noisy to obtain any meaningful quantification. However, there is a clear trend in the time series, with a calculated rate of 10 *µ*M*/*hour. This gives a sense of the limit of detection of the method.

**Figure 5:**
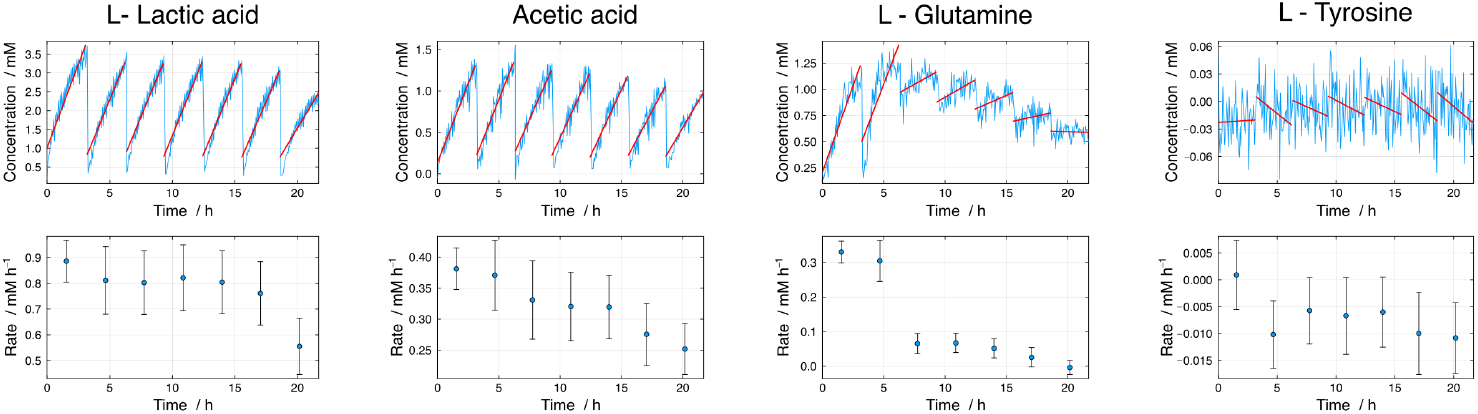
Comparison of time-resolved metabolic profiles and corresponding metabolic rates for selected metabolites.

Culture systems are increasingly used as biological models in the development of drugs, in particular to test drug candidates for efficacy and toxicity. This often involves challenge studies, where the behavior of the model system is compared in the presence and in the absence of the drug candidate in question. To demonstrate the principle of such a study with the TISuMR platform, we performed a comparative investigation using the well-established anti-psychotic drug Chlorpromazine, which is known to induce liver cholestasis in humans as a side effect.^62^

As a proof of principle, we conducted the same experiment described above; however, in this challenge study, the PCLS were treated with medium containing a mixture of human-like bile acid mixture (total bile-acid concentration was 16 *µ*M) and 20 *µ*M chlorpromazine. Small volumes of bile-acid stock solutions prepared in DMSO were added to the medium; the resulting incubation medium thus consisted of less than 0.5% DMSO. The concentration of chlorpromazine was specifically chosen due to its moderate toxicity for murine tissue slices incubated under static conditions in a 12-well plate; 20 *µ*M was close to the IC_50_ for this compound in the presence of bile acids.^63^ Under these conditions, we expected to be able to assess changed tissue responses compared to slices exposed to only the bile-acid mixture used as control.

Fig. 6 presents a comparison between metabolic rates of the control (blue circles) vs chlor-promazine-treated (green squares) precision-cut liver slices.

**Figure 6:**
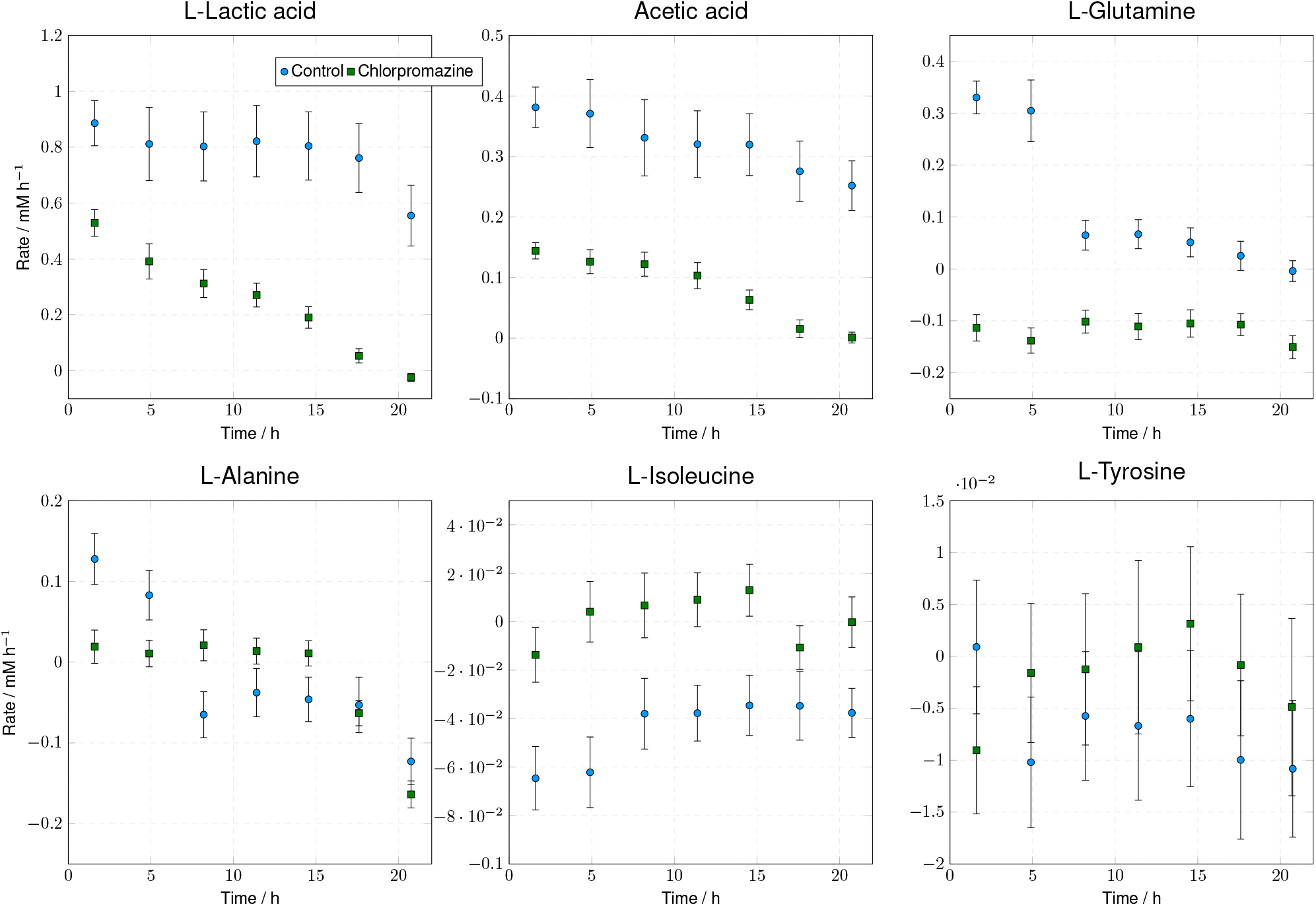
A comparison between metabolic rates of a control (blue circles) vs chlorpromazine-treated (green squares) precision-cut liver slices. Error bars are the 90% confidence interval of the fit.

The results reveal significant changes in metabolic profiles under drug treatment. For instance, under the control condition, L-lactic acid is produced at a rate of ~ 0.89mMh^−1^, while in the chlorpromazine-treated tissue, the initial production rate is lower (around ~ 0.5mMh^−1^) and progressively declines over time, eventually reaching zero. The rate of acetic acid production declined over time in both the control and challenge studies. However, the production rate in the control group was four times faster than in the group treated with chlorpromazine. L-glutamine is initially produced in the control (0.3 mMh^−1^) before stabilizing, while in treated tissue, it transitions to net consumption. Similarly, L-alanine remains stable or slightly positive in the control but shows clear negative rates following Chlorpromazine exposure. L-isoleucine is modestly consumed in both conditions, with slightly more pronounced uptake in treated samples. L-tyrosine shows low and variable rates near the method’s detection limit in both conditions, with a general trend toward uptake.

In a comparative study, Karsten *et al*.^63^ employed murine PCLS to evaluate the hepatotoxicity of chlorpromazine at varying concentrations. Notably, a 20 *µ*M dose was deemed moderately toxic after a 48-hour incubation in the presence of bile acids and Chlorpromazine but CPZ was only mildly toxic when no bile acids were present. Our TISuMR microfluidic perifusion platform demonstrated a more rapid decline in tissue metabolic output—approaching zero within just 24 hours under the same drug exposure. This accelerated response may be attributed to the platform’s continuos active perifusion, which ensures a more homogeneous drug distribution and dynamic nutrient exchange, potentially enhancing tissue-drug interaction. However, further validation is needed.

This proof-of-concept study demonstrates the significant potential of this platform to expand the range of metabolites that can be simultaneously monitored. However, several limitations must be addressed before it can be broadly adopted in biological research. One major drawback is that the current set up allows for the observation of only a single tissue slice at a time, which is not only cost-intensive due to the need for exclusive use of an NMR spectrometer throughout the observation period but also limits throughput. Consequently, obtaining statistically significant results from the same liver is currently impractical unless multiple dedicated spectrometers are available.

## Conclusions

The study demonstrates the TISuMR platform’s ability to detect and quantify metabolic changes in precision-cut liver tissue samples with high temporal resolution. The platform enables realtime monitoring of dynamic metabolic rates, providing detailed insights into metabolic activity and uncovering transient events that lower-resolution methods would miss.

The platform has the potential to be a powerful tool for investigating the metabolic impact of hepatotoxic drugs and in the future offer valuable potential for advancing drug safety studies and liver disease research. However, its current setup is limited to analyzing a single tissue slice at a time. This constraint reduces throughput and increases costs, limiting its practicality for large-scale studies. Addressing these limitations will be key to broadening the platform’s applicability and impact in biological research.

## Materials and Methods

### Microfluidic device fabrication

Microfluidic devices were fabricated from polycarbonate (PC) sheets (Weatherall Equipment, UK). The design of the device was created using AutoCAD 2016 and it comprised three layers: a bottom layer with a thickness of 175 *µ*m, a middle layer of 500 *µ*m, and a top layer of 250 *µ*m. Devices were cut out of PC sheet using a CO_2_ laser cutter (L3040 from HPC Laser LTD, Elland, UK). The bonding protocol firstly required a thorough cleaninng of each layer with isopropanol (Sigma-Aldrich, UK) followed by ethanol (Sigma-Aldrich), and a final rinse with isopropanol. After air drying, the bonding surfaces were treated with oxygen plasma for 80 seconds using a BD-20AC laboratory corona treater (Electro-Technic Products, USA). Then the layers were coated with 18 *µ*L of plasticiser (2.5 v/v % of dibutyl phthalate (Sigma-Aldrich) in isopropanol) and were placed in an oven at 65 °C for 15 minutes. Lastly the layers were bonded under 20 MPa of pressure at 85 °C for 15 minutes using a hydraulic press (Specac Ltd, Kent, United Kingdom). Each assembled microfluidic device was sterilized to use in perfusion experiments with PCLS.

### Tissue culture and Chlorpromazine treatment

The experimental workflow is depicted in Fig. 1a. Mouse liver tissue was collected from C57BL/6 mice aged 2 to 6 months. All procedures for obtaining the liver tissue were conducted in accordance with the Animal Act 1986 (scientific procedures) established by the UK Home Office. PCLS cores were prepared using a 6-mm biopsy punch from the mouse liver, which was kept in ice-cold University of Wisconsin (UW) solution (Bridge to Life, UK). The cores were sliced into PCLS using a Krumdieck Tissue Slicer (Alabama R&D, USA), maintaining them in ice-cold Krebs Henseleit Buffer 1x (pH 7.4). The PCLS were cultured for up to three days post-slicing, starting from day 0, in a 12-well plate with 1.3 ml of culture medium per well. The medium consisted of Williams Medium E (1X, glutaMAX-1) (Gibco, UK), supplemented with 0.13 g of D-Glucose (Merck, UK) for every 50 mL of culture medium, resulting in an overall D-Glucose concentration of 25 mM, along with 50 *µ*g/mL of Gentamicin (Gibco, UK) and 5 mM of deuterated HEPES buffer. The cultures were maintained in a tissue culture incubator at 37°C with an atmosphere of 80% oxygen and 5% CO_2_.

On the chip, the liver tissue-slice was placed in the central tissue culturing chamber. The delivery of nutrients and oxygen was provided by externally connected pumps and gas supply via the fluidic interface shown in Fig. 2a. First, a 25 *µ*L plug of Supplemented Williams Medium E was delivered into the channel marked in blue in Fig. 1b. A bidirectional pump was then used to pull the same plug of medium back-and-forth for 25 cycles. After that time, the plug was disposed of into a waste vial and a new plug of medium was introduced. This was achieved by pushing through 100 *µ*L of the culture medium. Carbogen was delivered into the adjacent gas channel (red) and its dissolution into the media channel was enabled by using a semi-permeable poly(dimethylsiloxane) PDMS membrane of 1 mm thickness over the top of the device. This ensured a continuous re-oxygenation of the supplied media. Lastly, physiological temperature of 37°C was maintained by equipping the fluidic interface with a PID controlled heating system as described elsewhere^30^ and illustrated in Fig. 2a. The flow of carbogen was controlled using a mass-flow controller placed at the end of the line while the media channel was back-pressurised to 1.2 bar in order to avoid bubble forming in the chip.

The composition of the bile acid mixture was obtained from Karsten *et al*. (2022) and it included 9.12 *µ*M cholic acid, 0.31 *µ*M chenodeoxy cholic acid, 2.87 *µ*M deoxycholic acid, 0.43 *µ*M hyodeoxycholic acid, 0.16 *µ*M lithocholic acid, 0.54 *µ*M ursodeoxycholic acid, 1.91 *µ*M taurocholic acid, 0.09 *µ*M taurochenodeoxycholic acid, 0.14 *µ*M taurohyodeoxycholic acid, 0.40 *µ*M sodium taurodeoxycholate hydrate, 0.03 *µ*M glycocholic acid, which resulted in 16 *µ*M bile acid (BA) mixture.^63^ All chemicals were purchased from Sigma-Aldrich (UK). In the control condition, on Day 1, a murine tissue was placed in the TISuMR device and was subjected to 16 *µ*M of human-like bile acid and DMSO mixture. The challenge study was performed by placing a tissue-slice into the TISuMR platform (on Day 1) and treating it with 20 *µ*M of Chlorpromazine + 16 *µ*M of bile acid mixture in the tissue culture media containing DMSO. This slice is monitored for up to 24 hours using the NMR.

### NMR experiments

The microfluidic device and two PDMS membranes were secured together using a 3D-printed fluidic interface as shown in Fig. 1a. This interface features liquid and gas in/out ports that allow for connections to external pumps and a gas cylinder for nutrient and oxygen delivery. The assembled device is then inserted into a custom-built NMR probe, which is equipped with a stripline micro-NMR detector optimized for the geometry of the 2.5 *µ*L sample detection chamber.

As illustrated in Fig. 1b, ^1^H NMR experiments were carried out on 14.1 T Bruker spectrometer, corresponding to a proton resonance frequency of 600 MHz, equipped with an Avance Neo console. Macro molecular contribution of the signal was suppressed using the T2 filter. Pure absorption multiplets were obtained by refocusing the J-coupling using the project sequence. A radio frequency pulse of 3.2 *µ*s (90° pulse) was used to acquire 8K points over a spectral width of 12 kHz (20 ppm). The strong signal present due to water is suppressed by continuous-wave presaturation for 2 s at the nutation frequency of 200 Hz. A data set of 48 transients was recorded with a repetition delay of 3 s. Free induction decay signals were Fourier transformed on 16K points with 1 Hz line broadening. Zero, first-order phase correction and baseline correction was applied to all the spectra. The chemical shift of all spectra was adjusted at D-Glucose doublets at 5.22 ppm. All the analysis of the obtained spectra was done using a program in Julia Language 1.6^64^ with NMR package written by Marcel Utz.^65^ Metabolite concentrations were obtained following the protocl in Ref.^30^ derived from NMR data contained in the human metabolome database.^61^

### ATP analysis

Following the NMR perfusion experiment, slices were retrieved from the TLP-PS probe within 10 minutes, placed in 1.5-mL Eppendorf tubes, snap-frozen in liquid nitrogen, and stored at −80°C.^66^ ATP content was later measured and normalized to protein concentration (ATP per mg protein) to assess metabolic activity. These values were compared to those of control PCLS maintained in a well plate under static incubation on the same day. In the ATP assay, the control PCLS showed an ATP content of 1.5 pmol/*µ*g protein, while slices treated with chlorpromazine exhibited no detectable ATP, indicating a complete loss of metabolic activity in the treated group.

## Acknowledgement

We gratefully acknowledge support by the H2020 FETOPEN project “TISuMR” (Grant number 737034).

## Credit Statement

SJB wrote the manuscript, created the figures, and analyzed and interpreted the experimental data. BP and MS conducted the experiments. RK was responsible for chlorpromazine sample preparation and contributed to the experimental design. EV and MU conceived the study and supervised the project. In addition, MU contributed to writing the manuscript.

